# Haplotype-resolved, chromosome-level assembly of white clover (*Trifolium repens* L., Fabaceae)

**DOI:** 10.1101/2023.06.06.543960

**Authors:** James S. Santangelo, Paul Battlay, Brandon T. Hendrickson, Wen-Hsi Kuo, Kenneth M. Olsen, Nicholas J. Kooyers, Marc T.J. Johnson, Kathryn A. Hodgins, Rob. W. Ness

## Abstract

**Background:** White clover (*Trifolium repens* L.; Fabaceae) is an important forage and cover crop in agricultural pastures around the world, and is increasingly used in evolutionary ecology and genetics to understand the genetic basis of adaptation. Historically, improvements in white clover breeding practices and assessments of genetic variation in nature have been hampered by a lack of high-quality genomic resources for this species, owing in part to its high heterozygosity and allotetraploid hybrid origin.

**Findings:** Here, we use PacBio HiFi and chromosome conformation capture (Omni-C) technologies to generate a chromosome-level, haplotype-resolved genome assembly for white clover totaling 998 Mbp (scaffold N50 = 59.3 Mbp) and 1 Gbp (scaffold N50 = 58.6 Mbp) for haplotypes 1 and 2, respectively, with each haplotype arranged into 16 chromosomes (8 per subgenome). We additionally provide a functionally annotated haploid mapping assembly (968 Mbp, scaffold N50 = 59.9 Mbp), which drastically improves on the existing reference assembly in both contiguity and assembly accuracy. We annotated 78,174 protein-coding genes, resulting in protein BUSCO completeness scores of 99.6% and 99.3% against the embryophyta_odb10 and fabales_odb10 lineage datasets, respectively.

**Conclusions:** We provide two white clover genome assemblies as part of this project: (1) a haplotype-resolved, chromosome-level assembly, and (2) a functionally annotated haploid mapping assembly. These assemblies place white clover among the best sequenced legumes to date, and one of the best assemblies for a plant of recent polyploid origins. This work promises to facilitate ongoing and future work in agricultural and evolutionary genetics in this agronomically and ecologically important species.

## Introduction

White clover (*Trifolium repens* L., Fabaceae) is a prostrate, herbaceous perennial that spreads via stolons, forming large clonal patches up to 1m across [1]. It originated as an allotetraploid in the Mediterranean 15 to 28 kya resulting from the hybridization of its diploid progenitors, *T. occidentale* and *T. pallescens* [2,3]. Because of its rapid growth and symbiosis with nitrogen-fixing bacteria, white clover is an important forage crop in agricultural pastures, and it has become naturalized in diverse climates around the world over the last several hundred years [1,4]. Today, there are large efforts to improve production and survival in variable environments, including traits such as yield and biomass production [5,6], salt tolerance [7], drought tolerance [8,9], frost tolerance [10,11], and disease resistance [12]. Currently, most white clover breeding relies on phenotypic selection, although marker-assisted breeding designs are increasingly common and would be greatly facilitated by a well-annotated, chromosome-level reference genome assembly [5,13].

In addition to its use in agricultural mixed-grass pastures and breeding programs, white clover has become a model in evolutionary ecology and genetics for understanding adaptation to environmental gradients and agents of selection in nature. Early work documented latitudinal and altitudinal clines in the frequency of cyanogenesis, the production of hydrogen cyanide (HCN) in response to tissue damage—an antiherbivore defense whose metabolic components can also affect tolerance to abiotic stressors (e.g., drought, frost) [14–18]. More recent work has corroborated these widespread continental clines [19–21], uncovered clines on smaller spatial scales across urban-rural gradients [22], identified the molecular mechanisms underlying genetic variation in cyanogenesis [23–26], and experimentally tested the ecological factors maintaining the cyanogenesis polymorphism [27–30] and its evolutionary consequences [31,32]. While much of this work has focused on the cyanogenesis polymorphism—a trait with well-characterized inheritance attributable to two Mendelian loci—ongoing and future work will leverage white clover’s rich history in evolutionary ecology to examine the genetic basis of adaptation at various spatial scales for which a high-quality reference assembly will be essential. In particular, a chromosome-level, haplotype-resolved assembly would facilitate identifying structural variants involved in adaptation [33] and improve our understanding of the evolutionary consequences of polyploidization in this ecologically and agronomically important allotetraploid.

Owing to the inherently repetitive nature of polyploid genomes, chromosome-level and haplotype-resolved genome assemblies have been challenging for these taxa. However, new technologies allow us to span difficult repetitive elements and offer the ability to greatly improve and expand earlier genome assemblies. Here we present a chromosome-level, haplotype-resolved genome assembly of the model legume white clover using PacBio HiFi and chromosome conformation capture (Hi-C) technologies. We present two genomes as part of this project: (1) an unannotated, haplotype-resolved assembly, and (2) a functionally annotated haploid mapping assembly, which we compare to the previous reference assembly [2] using two recently generated linkage maps for the species [23].

## Methods

### Plant sample

We sequenced an F4 *Trifolium repens* genotype that was generated as part of a separate experiment. This plant originated from an F0 crosses between a plant from Ontario, Canada, and a plant from Louisiana, USA, followed by two generations of random crossing. The sequenced plant was maintained in a 1L pot in potting soil (Pro-Mix LP15, Premier Tech, Rivière-du-Loup, Canada) in a growth chamber set to 25°C on a 12h light:12h dark cycle, though the plant was maintained in the dark for 48 hours prior to sampling to reduce polysaccharide content. The plant was non-destructively harvested on March 28, 2022, by sampling approximately 2.5g of leaf tissue, immediately flash-freezing tissue in liquid nitrogen, and storing it at –80°C prior to shipping on dry ice to Dovetail Genomics for DNA extraction, library preparation, and sequencing.

### Sequencing

DNA samples were quantified using Qubit 2.0 Fluorometer (Life Technologies, Carlsbad, CA, USA). The PacBio SMRTbell library (∼20 kbp mean insert length) for PacBio Sequel was constructed using SMRTbell Express Template Prep Kit 2.0 (PacBio, Menlo Park, CA, USA) using the manufacturer’s recommended protocol. The library was bound to polymerase using the Sequel II Binding Kit 2.0 (PacBio) and loaded onto PacBio Sequel II. Sequencing was performed on PacBio Sequel II 8M SMRT cells generating 58 Gbp of data. These PacBio CCS reads were used as an input to *hifiasm* v0.16.1-r375 [34,35] (see *Scaffolding* below).

For each Dovetail Omni-C library, chromatin was fixed in place with formaldehyde in the nucleus and then extracted. Fixed chromatin was digested with DNAse I, chromatin ends were repaired and ligated to a biotinylated bridge adapter followed by proximity ligation of adapter containing ends. After proximity ligation, crosslinks were reversed, and the DNA was purified. Purified DNA was treated to remove biotin that was not internal to ligated fragments. Sequencing libraries were generated using NEBNext Ultra enzymes and Illumina-compatible adapters. Biotin-containing fragments were isolated using streptavidin beads before PCR enrichment of each library. The library was sequenced on an Illumina HiSeqX platform (Illumina, San Diego, California, USA) to produce ∼30X sequence coverage. The PacBio CCS reads and Omni-C reads (MQ > 50) were then used as input for *hifiasm* to produce two haplotype-resolved assemblies (hap1 and hap2) using default parameters.

### Scaffolding

We first produced an initial assembly of all PacBio HiFi data with *hifiasm* in the “primary” mode. This resulted in two sets of contigs: primary and alternative. We then combined primary and alternative contigs into a single set of all contigs, containing 1,384,338,092 bp of sequence in 6,189 contigs with N50 size of 15,304,949 bp, which we call the “unresolved” contig set below. Next, to determine which contig was derived from which subgenome, we used Illumina reads for the diploid parental species *Trifolium occidentale* (SRR8593471) and *Trifolium pallescens* (SRR8617466) downloaded from NCBI SRA. We mapped the Illumina reads to the unresolved contig set with *bwa mem* [36], used the best alignment for each Illumina read, counted the number of reads from each parental species mapped to each contig, and divided it by the total number of Illumina reads in each parental set. Based on the weighted number of alignments of the parental reads, we labeled the contigs in the unresolved set with “Pall” and “Occ” labels corresponding to the two subgenomes, resulting in a “labeled” set of contigs.

We then aligned the PacBio HiFi reads and the Omni-C reads to the labeled contigs with the *minimap2* [37] and *bwa mem* aligners, computed the best alignment for each read, and split the HiFi and Omni-C reads into subsets for each subgenome. We required that both Omni-C reads have the best alignment to the same subgenome to be assigned to that subgenome. Next, we assembled the two subsets of HiFi/Omni-C reads separately with *hifiasm Hi-C* in haplotype resolved mode. This yielded haplotype-resolved assemblies for the two subgenomes. We then scaffolded the assemblies with *HiRise* scaffolder [38] and closed gaps in the scaffolds with *SAMBA* scaffolder [39]. The final step was to remove redundant haplotype contigs that *hifiasm* sometimes keeps in the assembly. We did this by aligning all contigs shorter than 1Mbp to the assembly for each of the haplotypes with *nucmer* [40] and excluding the contigs that mapped to the interior of other bigger contigs with better than 95% similarity over at least 75% of their length. This resulted in the final set of assembled haplotypes (2 subgenomes x 2 haplotypes each = 4 haplotypes).

### Assembly

All analyses from here forward are implemented in an open and reproducible Snakemake v7.16 pipeline [41]. The pipeline begins with input of the Dovetail haplotype assemblies, associated AGP (i.e., “A Golden Path”) files and linkage map data from [23], and ends with the generation of the phased diploid assembly in FASTA format (NCBI BioProjects PRJNA957817 and PRJNA957816), the annotated haploid mapping assembly in FASTA, NCBI Sequin, and GFF3 formats (BioProject PRJNA951196), and manuscript figures. See *Data Accessibility* for links to data and code.

Before assembling the reference genomes, the assembled haplotypes required manual curation to correct minor misassemblies and fill gaps generated during scaffolding. First, we used BLAST v2.12.0 [42] to align two previously generated linkage maps for the species [23] to each of the four haplotypes and to the previous *T. repens* reference genome [2]. We removed alignments that were less than 175 bp in length of the 200 bp total length for each linkage mapping marker sequence, and had less than 95% identity, and retained only the best alignment (i.e., lowest E-value) for each marker. These alignments were used to identify misassembled scaffolds and to assess correspondence between the scaffolds in the newly assembled haplotypes and the chromosomes in the previous reference genome.

Second, we used *Minimap2* v2.24 [37] to generate pairwise alignments between all four haplotypes. Together with the linkage map alignments above, these alignments enabled us to fill in three gaps (likely spanning the centromere) and one telomere with unplaced scaffolds (Figure S1). In addition, the scaffolding generated a double telomere at the end of one of the chromosomes in the *T. pallescens* subgenome; this extra telomere was removed and added to its correct location at the end of the homeologous chromosome in the *T. occidentale* subgenome (Figure S1). All manual fixes were implemented in BioPython v1.8 [43].

We used the revised haplotypes to generate two separate reference genome assemblies: a haplotype-resolved assembly and a collapsed haploid mapping assembly. As a diploidized allotetraploid, *T. repens* exhibits disomic inheritance with chromosomes from both subgenomes segregating independently. As such, its four haplotypes can be collapsed into two haplotypes, each containing 8 chromosomes from each subgenome (i.e., N = 16) resulting in a phased “diploid” assembly (i.e., 2N = 32). We therefore present this assembly as two FASTA files, with one for each of these two haplotypes. These FASTA files additionally include all unplaced scaffolds for each of the haplotypes. We additionally created a haploid mapping assembly, generated by taking the longer chromosome of each of the two haplotypes for each linkage group. This haploid mapping assembly was used for the structural and functional annotation described below. Both the diploid and haploid assembly were checked for annotation completeness by running BUSCO v5.4.6 [44] in “genome” mode against the embryophyta_odb10 and fabales_odb10 lineage datasets.

### Structural annotation and gene models

To improve gene-model predictions, we softmasked repeats prior to proceeding with annotation. First, we used *RepeatModeler* v2.0.3 [45] to generate a repeat library using the haploid mapping reference as input. This database was then merged with RepBase (v. RedBaseRepeatMaskerEdition-20181026), and the combined repeat library was used to softmask repeats using *RepeatMasker* v4.1.3.

We predicted gene models and generated a structural annotation of the haploid mapping assembly by combining evidence from proteins in related plant species and RNA-Seq evidence in *Trifolium repens*. First, we ran *BRAKER* v3.0.0 [46] in “protein mode” using proteins from all green plants (i.e., Viridiplantae) as input, supplemented with proteins from all legumes (Family: Fabaceae) from the UniProtKB database (N = 1,233,771 proteins). Next, we downloaded a subset of all RNA-Seq data from *T. repens* available from four published sources [2,10,47,48], selected to represent diverse tissue types and library preparation protocols (N = 21 RNAseq libraries total, Table S1). We mapped the raw RNA-Seq reads to the haploid mapping reference using *STAR* v2.7.0b [49] in “two-pass mode” and merged the resulting BAM files using SAMtools v1.16.1 [50]. We used this merged BAM file as input to *BRAKER* in “RNAseq” mode. Next, we combined evidence from “protein” and “RNAseq” modes using TSEBRA [51] with default parameters, except “intron_support” was reduced to 0.3 since the default was resulting in the loss of many transcripts. Finally, we used the *agat_convert_gxf2gxf*.*pl* script from *AGAT* v1.0.0–pl5321hdfd78af_0 [52] to convert the BRAKER-generated GTF to GFF3 format before proceeding to functional annotation.

### Functional annotation

We added functional annotations by querying extracted proteins against numerous databases prior to merging and formatting annotations for uploading to NCBI. First, we retrieved functional annotations using *InterProScan* v5.61.93-0 [53] and *Eggnog-mapper* v2.1.10 [54]. The resulting outputs were passed as input to *funannotate* v1.8.14 [55], which combined the annotations and queried some additional databases. For any protein annotated as “hypothetical protein” and containing a fully-resolved enzyme commission (i.e., EC) number (i.e., resolved to 4-digits), we replaced the “hypothetical protein” annotation with the EC number’s product in the ExPASSY Enzyme database [56]. If the EC number was only resolved to three digits or fewer, we kept the “hypothetical protein” annotation and removed the EC number. In the end, the following databases were queried for annotations: InterPro v93.0 [57], EggNog v5.0 [58], MEROPS v12.0 [59], Uniprot v2023_01 [60], dbCan v11.0 [61], Pfam v35.0 [62], GO v2023-03-06 [63,64], and MiBig v1.4 [65]. We queried our final annotation against the embryophyta_odb10 and fabales_odb10 BUSCO databases using BUSCO v5.4.6 [44] in “protein” mode and assessed synteny between the *T. occidentale* and *T. pallescens* subgenomes using *GENESPACE* with default parameters except “useHOGs” and “orthofinderInBlk” were set to TRUE [66].

## Results and Discussion

### Genome assemblies

Our final haplotype-resolved assembly totaled 998,247,995 bp for haplotype 1 (N = 693 scaffolds total) and 1,009,398,733 bp for haplotype 2 (N = 1,022 scaffolds total) (Table 1), slightly shorter than the ∼1.1 Gbp previously estimated genome size for *T. repens* species [2]. Haplotype 1 had genome BUSCO completeness scores of 99.6% and 99.5% against the embryophyta (N = 1,614 genes total) and fabales (N = 5,340 genes total) lineage datasets, respectively (Table 2). Similarly, haplotype 2 had BUSCO completeness scores of 99.5% against both databases.

**Table 1:** Assembly and annotation statistics for the haploid reference (16 chromosomes + 2 organelles) and the haplotype-resolved diploid reference assemblies.

| *Assembly statistics* |  |  |  |
| --- | --- | --- | --- |
|  | Reference |  |  |
|  | **Haploid** | **Diploid_Hap1** | **Diploid_Hap2** |
| **Number of contigs** | 18 | 693 | 1022 |
| **Largest contig** | 67,442,382 | 67,442,382 | 66,872,020 |
| **Total length** | 968,215,888 | 998,247,995 | 1,009,398,733 |
| **N50** | 59,895,999 | 59,287,556 | 58,622,306 |
| **N90** | 54,808,019 | 54,808,019 | 47,359,824 |
| **L50** | 8 | 8 | 9 |
| **L90** | 15 | 15 | 16 |
| **GC %** | 33.70 | 33.97 | 34.05 |
| **N's per 100 kbp** | 0.80 | 0.92 | 0.87 |
| *Annotation statistics* |  |  |  |
| **# genes** | 78,174 | — | — |
| **# genes with common names** | 15,871 | — | — |
| **# transcripts (mRNA)** | 87,929 | — | — |
| <u>Transcript level</u> |  |  |  |
| **Mean gene length** | 2,872 | — | — |
| **Total CDS exons** | 448,332 | — | — |
| **Single exon transcripts** | 18,658 | — | — |
| **Mean exon length** | 234 | — | — |
| **% of genome comprised of genes** | 23.2 |  |  |
| <u>Functional</u> |  |  |  |
| **GO terms** | 54,159 | — | — |
| **Interproscan** | 67,338 | — | — |
| **Eggnog** | 77,327 | — | — |
| **Pfam** | 55,891 | — | — |
| **Cazyme** | 3,313 | — | — |
| **Merops** | 3,039 | — | — |
| **Busco** | 3,408 | — | — |

**Table 2:** Genome BUSCO scores for all assemblies and Protein BUSCO scores for haploid mapping assembly against the embryophyta and fabales lineage datasets.

|  | *Genome-mode* |  | *Protein-mode* |  |
| --- | --- | --- | --- | --- |
|  | **Embryophyta_odb10 (N = 1,614)** | **Fabales_odb10 (N = 5,366)** | **Embryophyta_odb10 (N = 1,614)** | **Fabales_odb10 (N = 5,366)** |
|  | <u>Haploid</u> |  | <u>Haploid</u> |  |
| **Complete BUSCOs** | 99.6% (1,607) | 99.5% (5,340) | 99.6% (1,607) | 99.3% (5,328) |
| **Complete & single copy** | 5.3% (85) | 4.1% (221) | 3.9% (63) | 4.9% (264) |
| **Complete & duplicated** | 94.3% (1,522) | 95.4% (5,119) | 95.7% (1,544) | 94.4% (5,063) |
| **Fragmented** | 0.2% (3) | 0.1% (3) | 0.1% (2) | 0.1% (7) |
| **Missing** | 0.2% (4) | 0.4% (23) | 0.3% (5) | 0.6% (31) |
|  | <u>Haplotype 1</u> |  |  |  |
| **Complete BUSCOs** | 99.6% (1,608) | 99.5% (5,341) | – | – |
| **Complete & single copy** | 5.9% (95) | 4.1% (221) | – | – |
| **Complete & duplicated** | 93.7% (1,513) | 95.4% (5,120) | – | – |
| **Fragmented** | 0.2% (3) | 0.1% (3) | – | – |
| **Missing** | 0.2% (3) | 0.4% (22) | – | – |
|  | <u>Haplotype 2</u> |  |  |  |
| **Complete BUSCOs** | 99.5% (1,607) | 99.5% (5,339) | – | – |
| **Complete & single copy** | 5.6% (91) | 4.2% (225) | – | – |
| **Complete & duplicated** | 93.9% (1,516) | 95.3% (5,114) | – | – |
| **Fragmented** | 0.2% (4) | 0.1% (3) | – | – |
| **Missing** | 0.3% (3) | 0.4% (24) | – | – |

Our haploid mapping assembly totaled 968,215,888 bp assembled into 18 chromosomes (16 chromosomes + 2 organelles), with unplaced scaffolds removed from this assembly since our goal was to focus on assembled, contiguous chromosomes for this assembly. 93.8% (N = 2,186) of the 2,330 filtered linkage markers for the “DG” mapping population mapped to the correct chromosome in our new assembly, which was a dramatic improvement compared to the 39.4% (N = 919) of the correctly mapped markers from the previous assembly (Figure 1). Similar results were obtained for the “SG” mapping population (Figure S2). The haploid mapping assembly showed complete BUSCO scores of 99.6% and 99.5% against the embryophyta and fabales lineage datasets, respectively, with most of these genes occurring in duplicate copy (Table 2)

**Figure 1:**
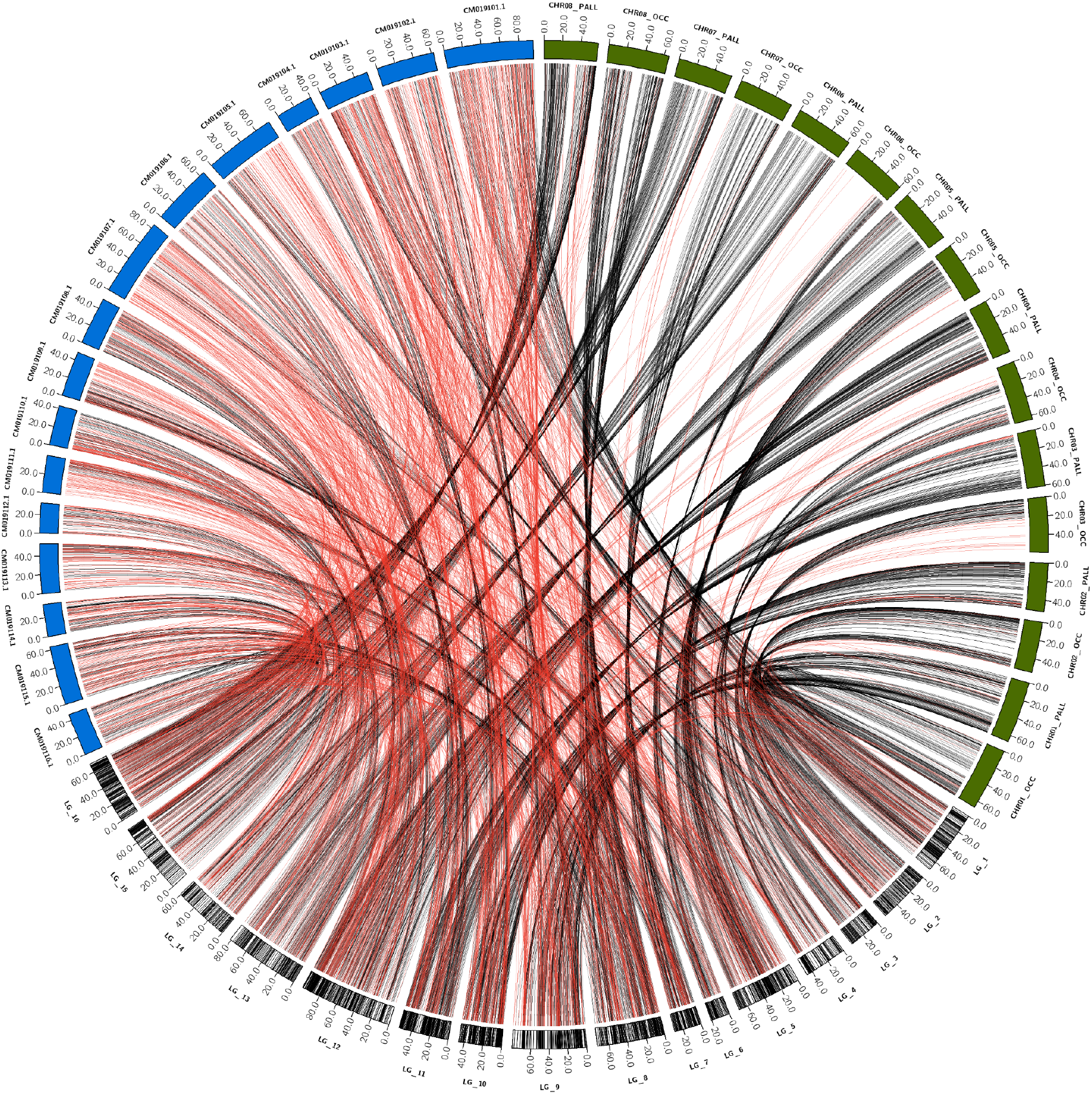
Linkage map from the “DG” mapping population ([23], bottom, converted to physical positions in Mbp) with markers (vertical black lines in ideogram) connected to their physical positions in both the previous reference assembly ([2], blue) and the current haploid assembly (green). Lines connecting markers to their physical position are colored red if they map to the wrong chromosome based on the linkage data, or black if they map to the correct chromosome. Similar figure for “SG” mapping population presented in supplementary figure S2.

**Figure 2:**
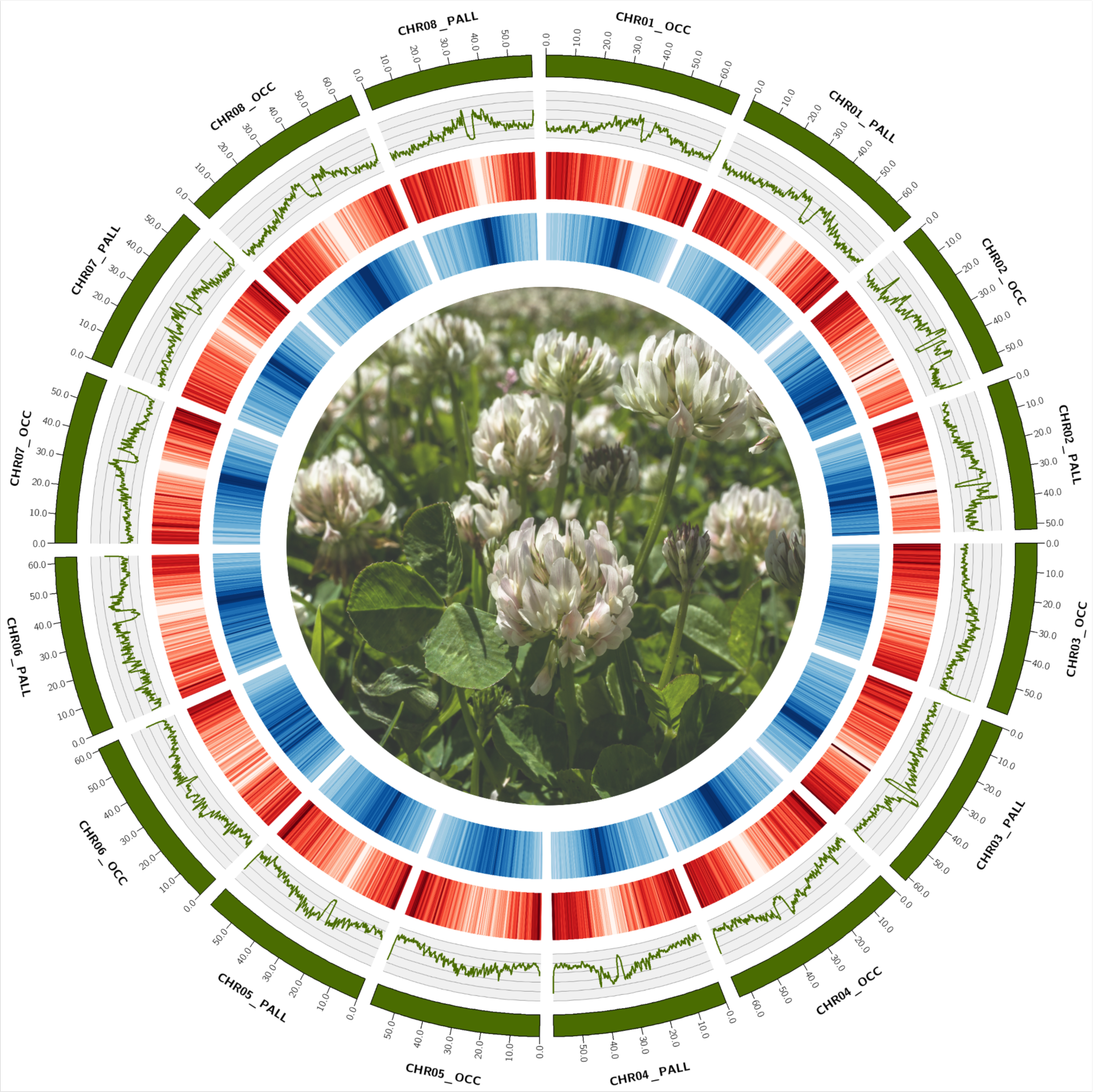
Circos plot of haploid mapping assembly consisting of 16 chromosomes. From outside to inside: chromosomes (green ideograms), GC%, gene density (red), repeat proportion (blue), and a photo of flowering *Trifolium repens* (credit: James Santangelo). GC%, gene density, and repeat density were estimated in 500 Kb windows with a 100 Kb step.

### Annotation

We softmasked 59.4% (∼576.5 Mbp) of the haploid reference assembly to improve gene model prediction during annotation. Of the classified repetitive elements, most (27.2%) were LTR elements, with Ty1/Copia (13.5%) and Gypsy/DIRS1 (9.7%) elements making up the majority (Table S2). We annotated 78,174 genes consisting of 87,929 mRNA transcripts that together account for 23.2% of the genome (Table 1). 39,425 of our annotated genes occur on the *T. occidentale* subgenome, with the remaining 38,749 on the *T. pallescens* subgenome, consistent with the number of genes of closely related diploid *Trifolium* species (*T. pratense:* 43,682; *T. subterraneum*: 42,704). Synteny between the subgenomes is largely preserved, except for 3 translocations between non-homeologous chromosomes and 6 inversions between homeologous chromosomes (Figure 3). Of the 78,174 genes, 4,868 (6.2%) are completely overlapped by repeats and likely represent transposable element protein-coding sequences. Most mRNAs (∼87%; N = 77,043) had at least one functional annotation (Table 1), with 15,871 genes containing common names. Our final annotated protein set had complete protein BUSCO scores of 99.6% and 99.3% against the embryophyta and fabales lineage datasets, respectively.

**Figure 3:**
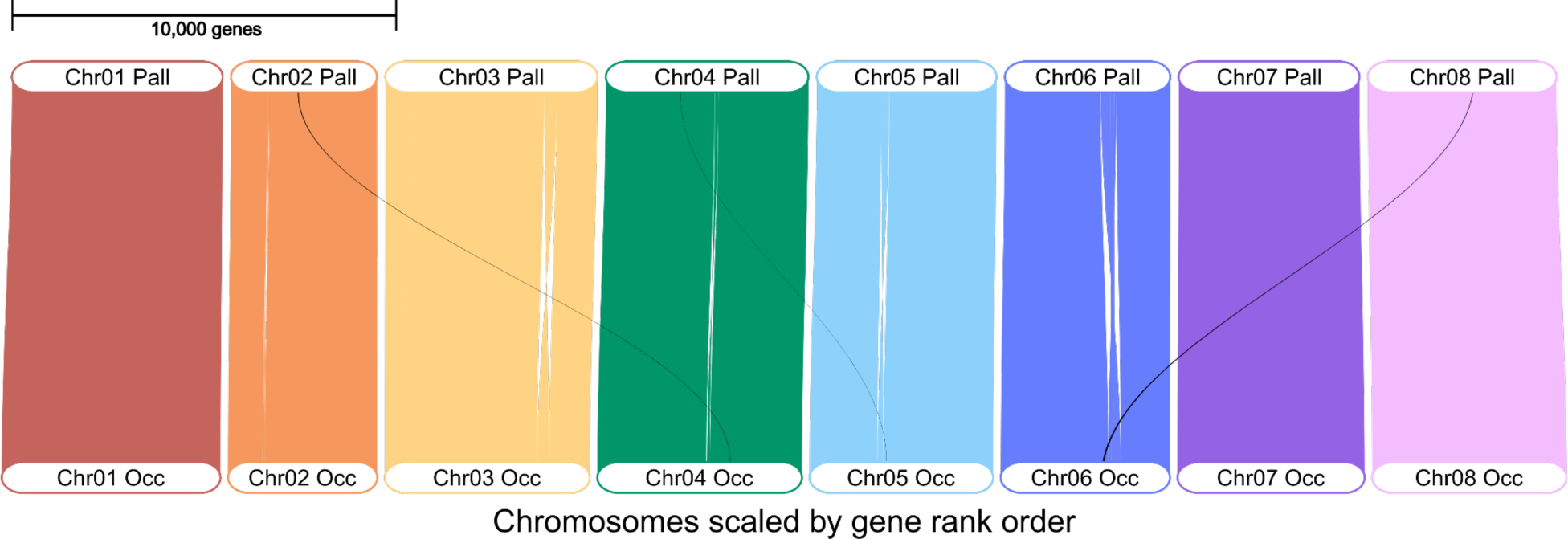
GENESPACE Riparian plot showing synteny between the *T. occidentale* (bottom) and *T. pallescens* (top) subgenomes of *T. repens*. Black lines show inferred translocations between non-homeologous chromosomes (N = 3), while white gaps within homeologous chromosomes show inversions (N = 6; Chr_06 contains two nested inversions).

## Conclusion

We have provided a chromosome-level, haplotype-resolved genome assembly of the allotetraploid white clover (*Trifolium repens*), and a functionally annotated haploid mapping assembly that shows substantial improvements over the existing reference genome for the species. These assemblies will facilitate marker-assisted breeding programs for traits of agronomic importance and provide increased resolution and ability to identify the genomic basis of adaptation in this increasingly used model in evolutionary ecology and genetics. Together with an alternative and upcoming chromosome-level assembly [67] and other high-quality reference genomes in the genus [68–70], our haplotype-resolved assembly will be particularly useful for identifying structural variation and facilitate the development of pangenomic references [71] for which haplotype-resolved assemblies are an asset [72].

## Data Accessibility

All code to reproduce this manuscript’s results can be found on JSS’s GitHub (see https://github.com/James-S-Santangelo/dcg). The pipeline requires the raw haplotypes assembled by Dovetail and the AGP files, which are currently self-hosted (https://ln5.sync.com/dl/ae3e80d10/822i24ai-uhhhg646-k6u7d8ha-qsmm4wc3) and will be permanently archived following publication. All other data is contained within the GitHub repository, except for proprietary databases (e.g., RepBase) that could not be included (see GitHub README). The NCBI BioProjects containing the assemblies and annotation files are currently public but are in the final stages of processing and may not be available yet. In the meantime, NCBI-approved assemblies and annotation files can be found here: https://ln5.sync.com/dl/9ef9902a0/x87rgt8f-za3ttvse-nj8bd3va-yvaews78.

## Author Contributions

JSS: Conceptualization, sample preparation, software, formal analysis, investigation, data curation, writing—original draft, writing—review and editing, visualization.

PB: Software, formal analysis, writing—review and editing.

BTH: Software, formal analysis, data curation—review and editing.

WHK: Validation, writing – review and editing.

KMO: Writing – review and editing.

NJK: Conceptualization, writing—review and editing, funding acquisition.

MTJJ: Conceptualization, sample preparation, interpretation, reviewing and editing, funding acquisition.

KAH: Conceptualization, writing—review and editing, funding acquisition.

RWN: Conceptualization, writing—review and editing, funding acquisition.

## Acknowledgements

We thank Rory Craig for thoughtful discussions and comments on an early draft of the manuscript. Plant lines used for sequencing were originally created by L. Albano, and subsequently maintained by H. Fargo, K. Bhachu, and I. Arif. DNA extraction, genome sequencing and scaffolding was provided by Dovetail genomics.

## Supplemental materials

**Figure S1:**
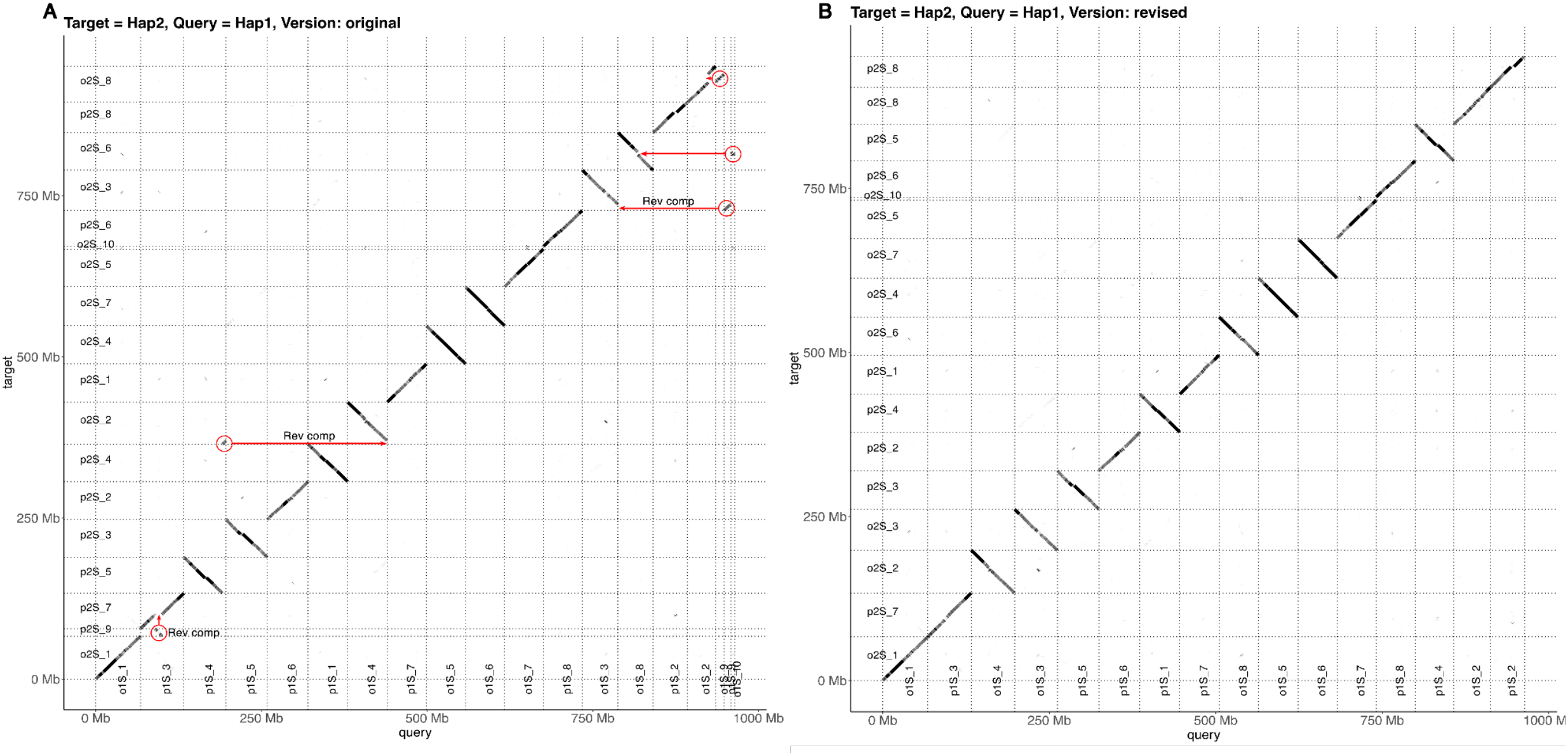
Original Dovetail haplotypes (A) and revised haplotypes (B) following manual fixes implemented in BioPython. Red circles surround fragments (often entire unplaced scaffolds) that map to gaps or telomeric regions of assembled chromosomes. Red arrows indicate where fragments were placed, and whether they were reverse-complemented before placement.

**Figure S2:**
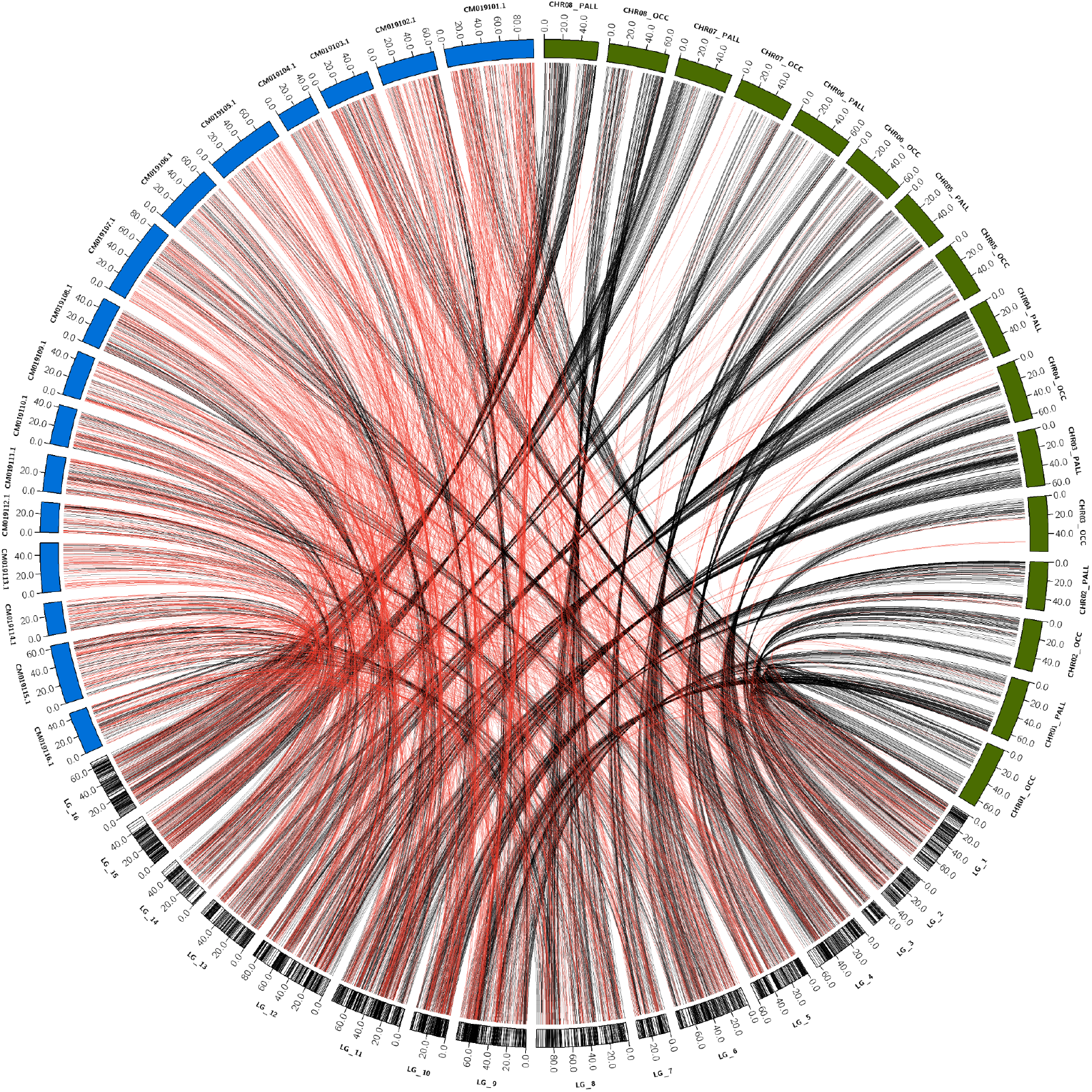
Linkage map from the “SG” mapping population ([23], bottom, converted to physical positions in Mbp) with markers (vertical black lines in ideogram) connected to their physical positions in both the previous reference assembly (blue) and the current haploid assembly (green). Lines connecting markers to their physical position are colored red if they map to the wrong chromosome based on the linkage data, or black if they map to the correct chromosome. 94.6% (N = 2,049) of the 2,165 filtered linkage markers mapped to the correct chromosome in our new assembly, compared to 40.0% (N = 867) in the previous assembly.

## Supplementary tables

**Table S1:** RNAseq accessions used for genome annotation using BRAKER.

| Accession | Tissue | # Bases (Gbp) | Selection | Reference |
| --- | --- | --- | --- | --- |
| SRR12578192 | Root | 6.9 | oligot-dT | [47] |
| SRR12578220 | Root | 7.8 | oligot-dT | [47] |
| SRR12578237 | Root | 7.6 | oligot-dT | [47] |
| SRR12578285 | Root | 10.2 | oligot-dT | [47] |
| SRR12578290 | Root | 8.2 | oligot-dT | [47] |
| SRR8691037 | Root | 19.5 | cDNA | [2] |
| SRR8691038 | Stolon | 17 | cDNA | [2] |
| SRR8691039 | Leaf | 17.7 | cDNA | [2] |
| SRR8691040 | Flowers | 18.7 | cDNA | [2] |
| SRR21383273 | Root | 5 | Random | [73] |
| SRR21383275 | Root | 4.8 | Random | [73] |
| SRR21383276 | Root parasitized by<br>Cuscuta | 4.6 | Random | [73] |
| SRR21383278 | Root parasitized by<br>Cuscuta | 4.7 | Random | [73] |
| SRR20706441 | Root | 5 | Random | [73] |
| SRR20706444 | Root | 4.8 | Random | [73] |
| SRR5814845 | Flowers | 5.4 | cDNA | [74] |
| SRR5814846 | Flowers | 6.6 | cDNA | [74] |
| SRR5814847 | Flowers | 6.3 | cDNA | [74] |
| SRR5814848 | Flowers | 7.3 | cDNA | [74] |
| SRR5814849 | Flowers | 6.2 | cDNA | [74] |
| SRR5814850 | Flowers | 6.1 | cDNA | [74] |

**Table S2:**
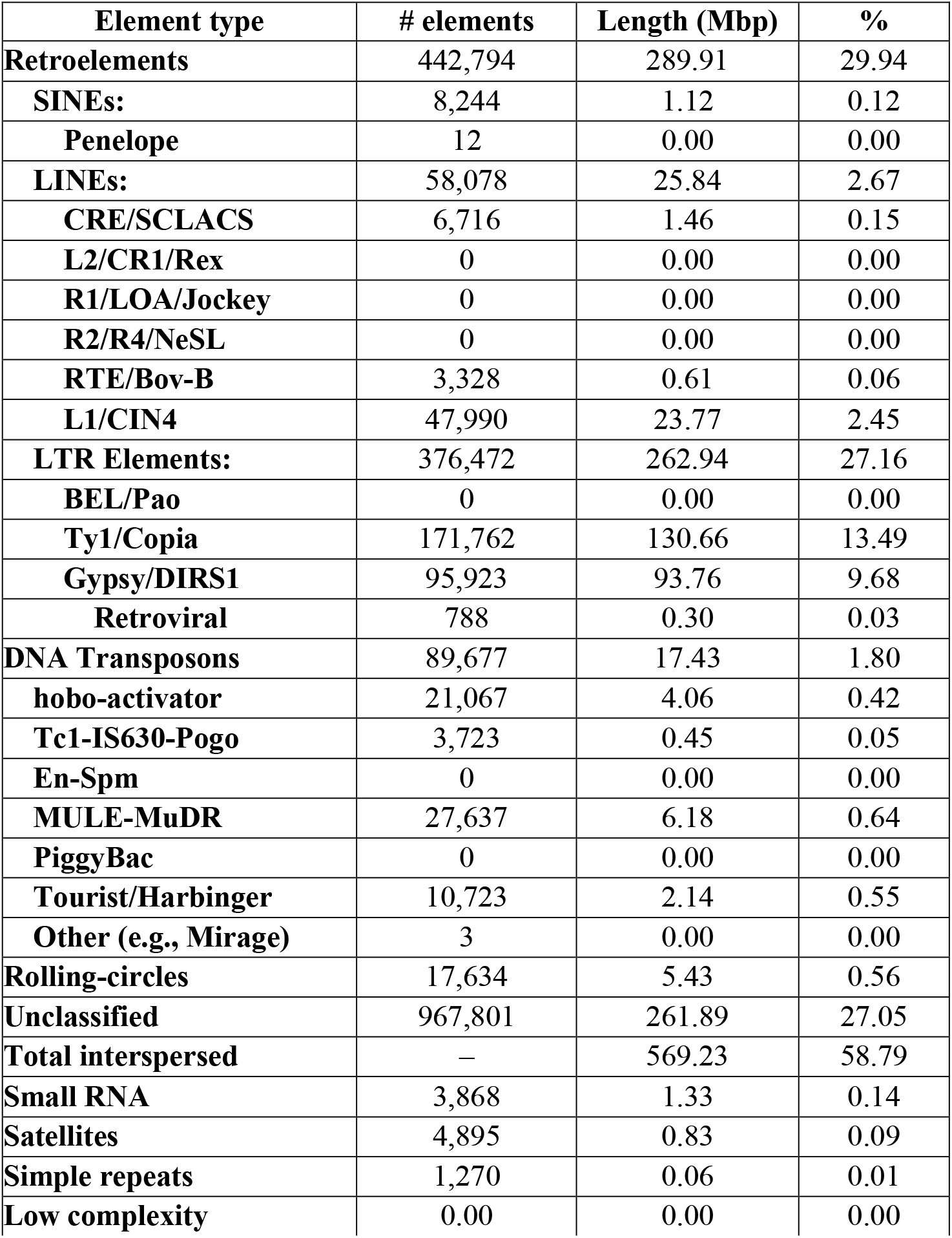
Type, number, length, and proportion of repeat elements in the *T. repens* haploid mapping assembly based on Repeat Modeler and Repeat Masker analysis.

| **Element type** | **# elements** | **Length (Mbp)** | **%** |
| --- | --- | --- | --- |
| **Retroelements** | 442,794 | 289.91 | 29.94 |
| **SINEs:** | 8,244 | 1.12 | 0.12 |
| **Penelope** | 12 | 0.00 | 0.00 |
| **LINEs:** | 58,078 | 25.84 | 2.67 |
| **CRE/SCLACS** | 6,716 | 1.46 | 0.15 |
| **L2/CR1/Rex** | 0 | 0.00 | 0.00 |
| **R1/LOA/Jockey** | 0 | 0.00 | 0.00 |
| **R2/R4/NeSL** | 0 | 0.00 | 0.00 |
| **RTE/Bov-B** | 3,328 | 0.61 | 0.06 |
| **L1/CIN4** | 47,990 | 23.77 | 2.45 |
| **LTR Elements:** | 376,472 | 262.94 | 27.16 |
| **BEL/Pao** | 0 | 0.00 | 0.00 |
| **Ty1/Copia** | 171,762 | 130.66 | 13.49 |
| **Gypsy/DIRS1** | 95,923 | 93.76 | 9.68 |
| **Retroviral** | 788 | 0.30 | 0.03 |
| **DNA Transposons** | 89,677 | 17.43 | 1.80 |
| **hobo-activator** | 21,067 | 4.06 | 0.42 |
| **Tc1-IS630-Pogo** | 3,723 | 0.45 | 0.05 |
| **En-Spm** | 0 | 0.00 | 0.00 |
| **MULE-MuDR** | 27,637 | 6.18 | 0.64 |
| **PiggyBac** | 0 | 0.00 | 0.00 |
| **Tourist/Harbinger** | 10,723 | 2.14 | 0.55 |
| **Other (e.g., Mirage)** | 3 | 0.00 | 0.00 |
| **Rolling-circles** | 17,634 | 5.43 | 0.56 |
| **Unclassified** | 967,801 | 261.89 | 27.05 |
| **Total interspersed** | — | 569.23 | 58.79 |
| **Small RNA** | 3,868 | 1.33 | 0.14 |
| **Satellites** | 4,895 | 0.83 | 0.09 |
| **Simple repeats** | 1,270 | 0.06 | 0.01 |
| **Low complexity** | 0.00 | 0.00 | 0.00 |

